# Fitness landscape analysis of a tRNA gene reveals that the wild type allele is sub-optimal, yet mutationally robust

**DOI:** 10.1101/2021.09.27.461914

**Authors:** Tzahi Gabzi, Yitzhak Pilpel, Tamar Friedlander

**Affiliations:** Department of Molecular Genetics, Weizmann Institute of Science, Rehovot 7610001, Israel; The Robert H. Smith Institute of Plant Sciences and Genetics in Agriculture Faculty of Agriculture, Hebrew University of Jerusalem, 229 Herzl St., Rehovot 7610001, Israel

## Abstract

Fitness landscape mapping and the prediction of evolutionary trajectories on these landscapes are major tasks in evolutionary biology research. Evolutionary dynamics is tightly linked to the landscape topography, but this relation is not straightforward. Here, we analyze a fitness landscape of a yeast tRNA gene, previously measured under four different conditions. We find that the wild type allele is sub-optimal, and 8%-10% of its variants are fitter. We rule out the possibilities that the wild type is fittest on average on multiple conditions or located on a local fitness maximum. Instead, we find that the wild type is mutationally robust (‘flat’), while more fit variants are typically mutationally fragile. Similar observations of mutational robustness or flatness have been so far made in very few cases, predominantly in viral genomes.

## Introduction

Fitness landscape mapping and prediction of evolutionary trajectories on these landscapes are major tasks in evolutionary biology [1]. While evolutionary theory predicts that population mean fitness should increase over time, it offers only few quantitative predictions for the dynamics of evolution and the possible evolutionary trajectories. The main hurdle for generally computing evolutionary trajectories is their dependence on the underlying fitness landscape. Currently available fitness landscapes include between 16 and 100,000 different genotypes (for review see [2, 3]). Yet, even the largest datasets [4, 5, 6, 7] encompass only small fractions of the entire fitness landscape of even a single gene. As detailed fitness measurements have been unavailable until recently, most of the associated theory was developed in isolation from data [8, 9, 10, 11, 12, 13, 14, 15, 16].

The advent in sequencing technologies now enables measurement of increasingly larger fitness landscape datasets [6, 7]. It is then desirable to predict evolutionary trajectories on these empirical fitness landscapes, using the previously developed theory in this field.

A recent set of experiments characterized the fitness landscape of the tRNA^Arg^ccu gene of *S. cerevisiae*. As this gene is relatively short (72 nucleotides), its landscape is significantly smaller than that of a typical protein. It is a single-copy, non-essential gene, such that many of its mutants are viable. Li *et al*. measured the growth rates of 23, 284 different variants of this gene (Fig. 1a) under four different growth conditions (23°C, 30°C, 37°C and oxidative stress) that represent well typical growth conditions during life of yeast in nature [17, 18]. The richness of this dataset renders it a highly valuable case study for analyzing topographic properties and evolutionary trajectories of an empirical fitness landscape and for comparing them with theoretical predictions. In analyzing this fitness landscape we noticed that many variants appear fitter than the wild-type in each of the examined conditions. The wild-type’s advantage appears instead to be in its mutational robustness, since its neighbors in sequence space are relatively fit too.

**Figure 1:**
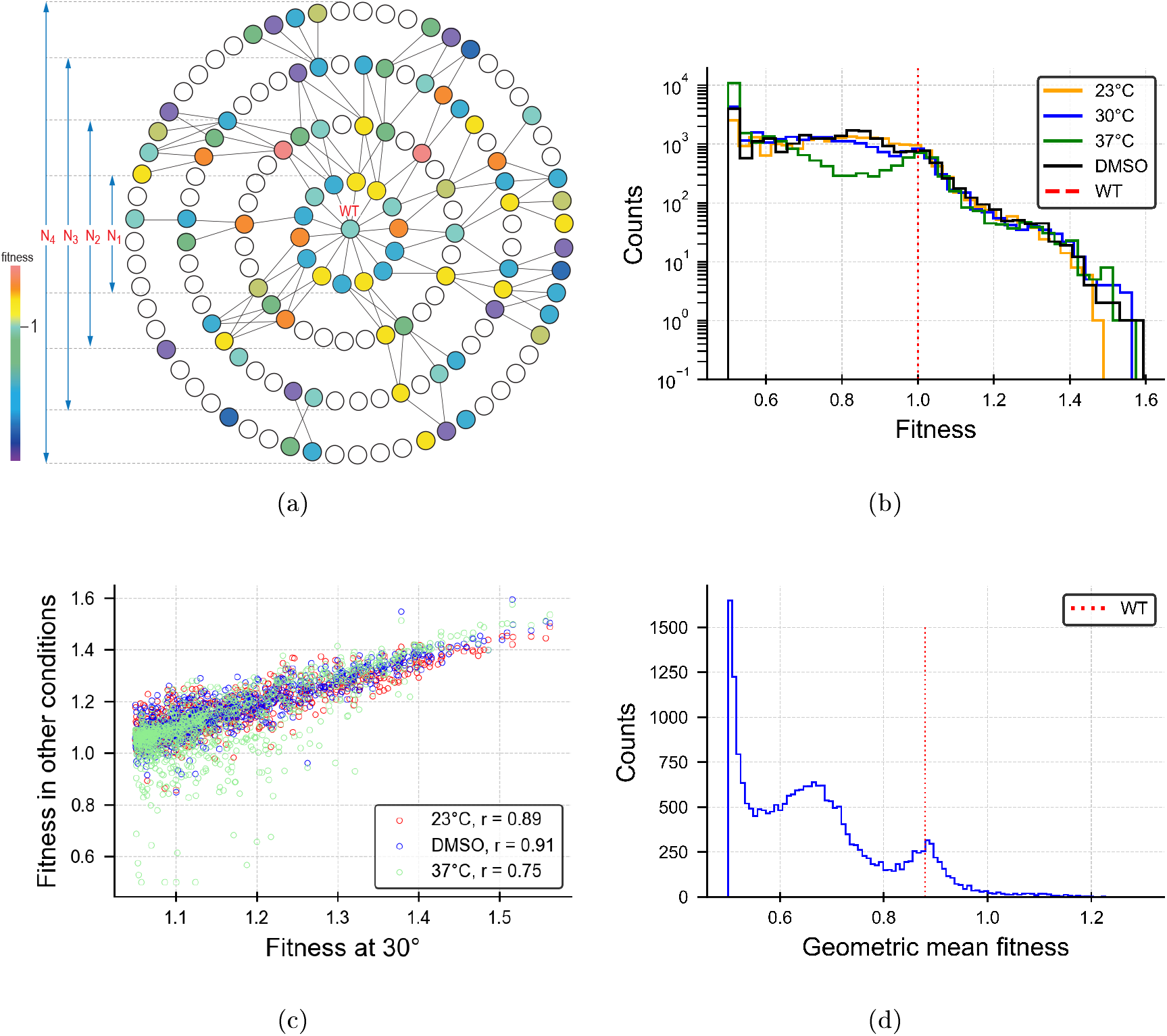
The wild type is not the fittest at any of the conditions or on average on all four. **(a)** A schematic visualization of the experimentally measured tRNA fitness landscape. Each circle represents a genotype. Filled circles represent genotypes whose fitness values (here encoded by different colors) were measured. Empty circles represent genotypes whose fitness values were not measured. We use here a concentric representation of the fitness landscape, centered around the wild type, where the minimal number of steps on the graph between any two genotypes is the number of point mutations separating them. The wild type is then surrounded by expanding circles of its single mutants (denoted by *N*_1_, double mutants (*N*_2_, etc. The experiment probed all the wild type’s single-point mutants, but only decreasingly smaller proportions of the following mutational neighborhoods, *N_i_*. **(b)** The distribution of all fitness values measured under four different conditions (23°C, 30°C, 37°C and DMSO), at semi-log scale. The wild type fitness value is shown by the red dotted line. Fitness was defined relative to the wild type’s fitness, such that the wild type fitness was set to 1 for each condition. Under each of the conditions tested, 8%-10% of the genotypes in this dataset were fitter than the wild type, **(c)** Fitness values of variants with fitness in the range [1.05, 1.6] at 30°C plotted against their fitness values at 23°C, DMSO and 37°C. The correlation coefficient between fitness values under different conditions were *r* = 0.89, 0.91, 0.75 respectively. We conclude that variants that have high fitness in one environment usually have high fitness in all four of them. **(d)** Distribution of 〈*f_i_*〉 the geometric mean fitness values over the four environments for all genotypes. Note that after alignment of fitness values under different conditions to a common baseline, the wild type fitness is no longer 1 (Methods).

## Results

A remarkable observation is that the wild type is not the genotype with highest fitness under any of the four conditions. Under each of the conditions, between 8%-10% of the variants exhibited higher fitness than the wild type (Fig. 1b) We then analyzed possible sources for measurement errors, including statistical sampling fluctuations in read-counts, as a source of inaccuracy in fitness assessment and the possibility that the fitness effect was due to independent mutations that fortuitously occurred elsewhere in the genome (SI). We conclude that although such errors do exist, they could not fully account for the wild type’s fitness sub-optimality.

A possible explanation for the apparent sub-optimality of the wild type could be that while some variants are fitter than the wild type under a specific condition, they are much less fit under other conditions, such that, *on average* across conditions the wild type is fittest. To test this explanation, we checked for all high-fitness genotypes (*f*>1.05 at 30°C) the correlation between their fitness values under the various growth conditions - Fig. 1c. We found, that most genotypes which are fit under one condition are also fit under others.

To formally compare between fitness values averaged over multiple conditions, we also calculated the geometric mean fitness [19], 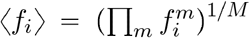, where 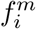 is the fitness value of the *i*-th genotype in the *m*-th condition out of *M* (Methods).

Fig. 1d shows a histogram of the geometric mean fitness values 〈*f_i_*〉 of all the genotypes in our dataset. Here too, we observe that the wild type is not the fittest across conditions.

Both results argue against the possibility that the wild type is the fittest on average across conditions. A possible caveat is that only four conditions were included in this calculation, and some of the high-fitness mutants could be inferior in another condition not included in this experiment. However, the conditions measured in the experiment are representative of yeast natural habitat and the wild type is sub-optimal in several of the most highly relevant conditions. Additionally, a recent study that analyzed fitness of yeast in 45 different growth conditions found correlations in fitness between conditions [20], thus arguing against the possibility that many genotypes behave very differently in some other condition(s).

Alternatively, the wild type sub-optimality could hypothetically be rooted in the fitness landscape topography. If, for example, the wild type were an isolated local maximum, separated from the global fitness maximum by fitness valleys, the population could be “trapped” in the current wild type genotype, hindered from reaching the global maximum (at least temporarily) [8, 9]. To test this hypothesis, we started by locating the high-fitness genotypes in the dataset. We define “Mutational neighborhoods” *N_i_*(WT) surrounding the wild type as the set of genotypes reachable by *i* point mutations (shortest path) from the wild type (see Fig. 1a). Fig. 2a shows the fitness distributions of the four mutational neighborhoods *N*_1_ – *N*_4_ (single to quadruple mutants). We found that all four mutational neighborhoods contained fitter-than wild type genotypes, but the largest proportion of such fitter genotypes was in *N*_2_, only two point mutations away from the wild type.

**Figure 2:**
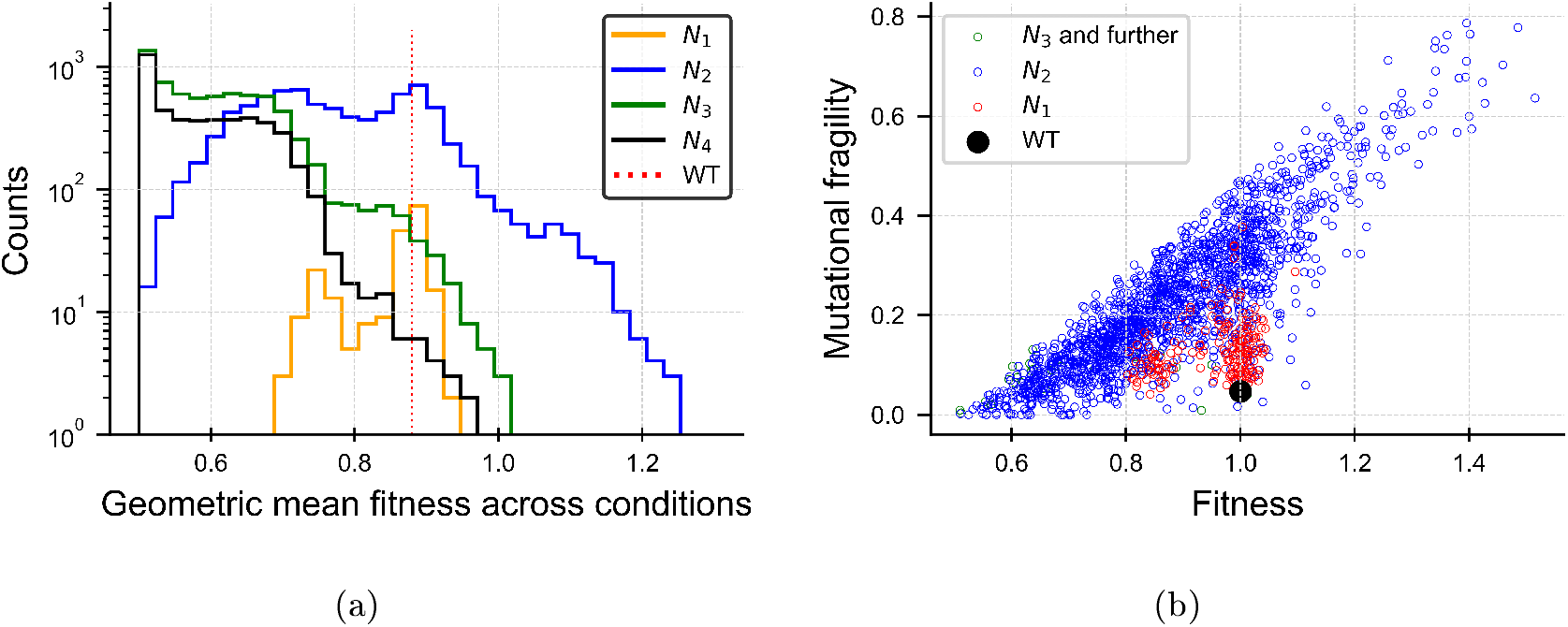
The wild type is not a local maximum and is insensitive to mutations. **(a)** Fitness value histograms of genotypes in the wild type’s four mutational neighborhoods *N*_1_ – *N*_4_. The wild type fitness (red dotted line) is shown for reference. Notably, all four mutational neighborhoods contain fitter than wild type genotypes, but the largest proportion of fitter genotypes is in *N*_2_. **(b)** Genotype mutational fragility against fitness - scatter plot. The wild type (black) is nearly the least fragile to deleterious mutations amongst all genotypes with similar fitness values. Low-fitness genotypes often have mostly beneficial mutations and hence their fragility is low. We show here all genotypes with at least 5 nearest-neighbors, such that at least 3 of them are further away from the wild type (see Methods). We illustrate genotypes that are wild type single, double or higher mutants using different colors.

Dissection of each mutational neighborhood into one of three fitness categories shows that 69% (142 out of 207) of the wild type’s single mutants have similar geometric mean fitness values to the wild type’s (|〈*f_i_*〉 – 〈*f*(WT)〉| < Δ), with Δ = 0.05, 29% (61 genotypes) of them were much less fit (〈*f_i_*〉 < 〈*f*(WT)〉 – Δ), and 2% (4 genotypes) were fitter than the wild type by more than Δ = 0.05, (〈*f_i_*〉 > 〈*f*(WT)〉 + Δ). Amongst the *N*_2_(WT) genotypes (wild type’s double mutants) the proportion of such fitter-than wild type genotypes (〈*f_i_*〉 > *f*(WT) + Δ) was even larger (730 out of 8101; 9%).

We then checked for the existence of evolutionary trajectories of non-decreasing fitness, leading from the wild type to the fitter genotypes in *N*_2_ and *N*_3_. We mapped all 2- and 3-step trajectories of strictly increasing geometric mean fitness (or fitness at 30°C, numbers in brackets), originating from the wild type. The number of fitness increasing trajectories depends on the minimal required fitness difference in both steps. For example, if we require at least 1% fitness increase at each such step, we find 277 (480) length-3 and 771 (1024) length-2 possible trajectories starting from the wild type. If we require at least 2% fitness increase at each step, the number of trajectories decreases, but there are still 26 (133) length-3 and 159 (425) length-2 trajectories. By requiring a minimal fitness increase, we ensure that such trajectories are strictly increasing, and are not mistakenly classified as such due to inaccuracies in fitness estimations. This is certainly an under-estimation of the number of fitness-increasing trajectories, because evolutionary transitions to genotypes with smaller fitness advantage or even to neutral ones are also possible and were not included in this enumeration. Additionally, the dataset we used does not include all double and triple mutants, which could also add to the trajectory count.

We conclude that the wild type is not a local maximum which is mutationally isolated from higher-fitness genotypes. It is worth mentioning in this context that the higher the landscape dimension is, the larger the number of possible single mutants for each genotype is. A genotype is only a local maximum if *all* its single mutants are less fit. Hence, with the increase of landscape dimensionality, it is less likely to find local maxima [3].

Up to this point, we found that the wild type is neither the fittest on average, nor is it a local maximum. What is then unique about this genotype and why was this sequence selected in evolution to be the wild type? In Fig. 2a we saw already that most of the wild type nearest-neighbors are similar in fitness to the wild type. Hence, we next sought to characterize whether such fitness “flatness” is common in this fitness landscape or whether the wild type is unique in residing in a relatively flat region of the landscape. We defined genotype mutational fragility as the average fitness difference between the genotype and its deleterious single-mutants (beneficial single-mutants receive zero weight):

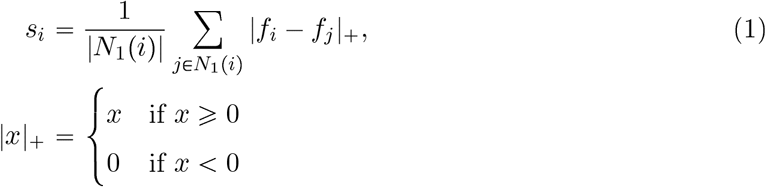

where |*N*_1_(*i*)| is the total number of single mutants of genotype *i*. This definition is based on fitness information of all the single mutants of genotype *i*. In practice, with the exception of the wild type, we only had measurements of a subset of the single mutants and estimated *s_i_* using partial data. To minimize biases because of small numbers of single mutants, fragility was only calculated for genotypes having at least 5 single-mutants, 3 of which are further away from the wild type (Methods). This limitation enabled calculation only for 1854 variants out of 23,284.

Fig. 2b shows a scatter plot of genotype mutational fragility plotted against genotype fitness. Interestingly, we observe that the wild type is at the ‘tip’ of the fragility-fitness cloud, such that it is nearly the least fragile amongst genotypes with similar fitness value and nearly the fittest amongst genotypes with similarly low fragility. Hence, it exhibits a balance between fitness and mutational robustness. What is the relation between fragility and fitness in other landscapes? We used the NK model [13] to test this question. Here we show simulation results with parameter values *N* = 14, *K* = 5,13 (Figs. 3a-b). In this example, the fitness values of all genotypes are known and fragility can be calculated with no sampling bias. *K* = *N* – 1 = 13 represents an uncorrelated landscape and *K* = 5 represents a partially correlated one. In both cases we found a similar crescent-shaped fitness-fragility relation, reminiscent of the relation we obtained in the tRNA dataset. The differences in fragility between the intermediate and the extreme fitness genotypes were larger in the uncorrelated landscape (Fig. 3b) than in the partially-correlated one (Fig. 3a), while their fitness values distributions were similar (SI).

**Figure 3:**
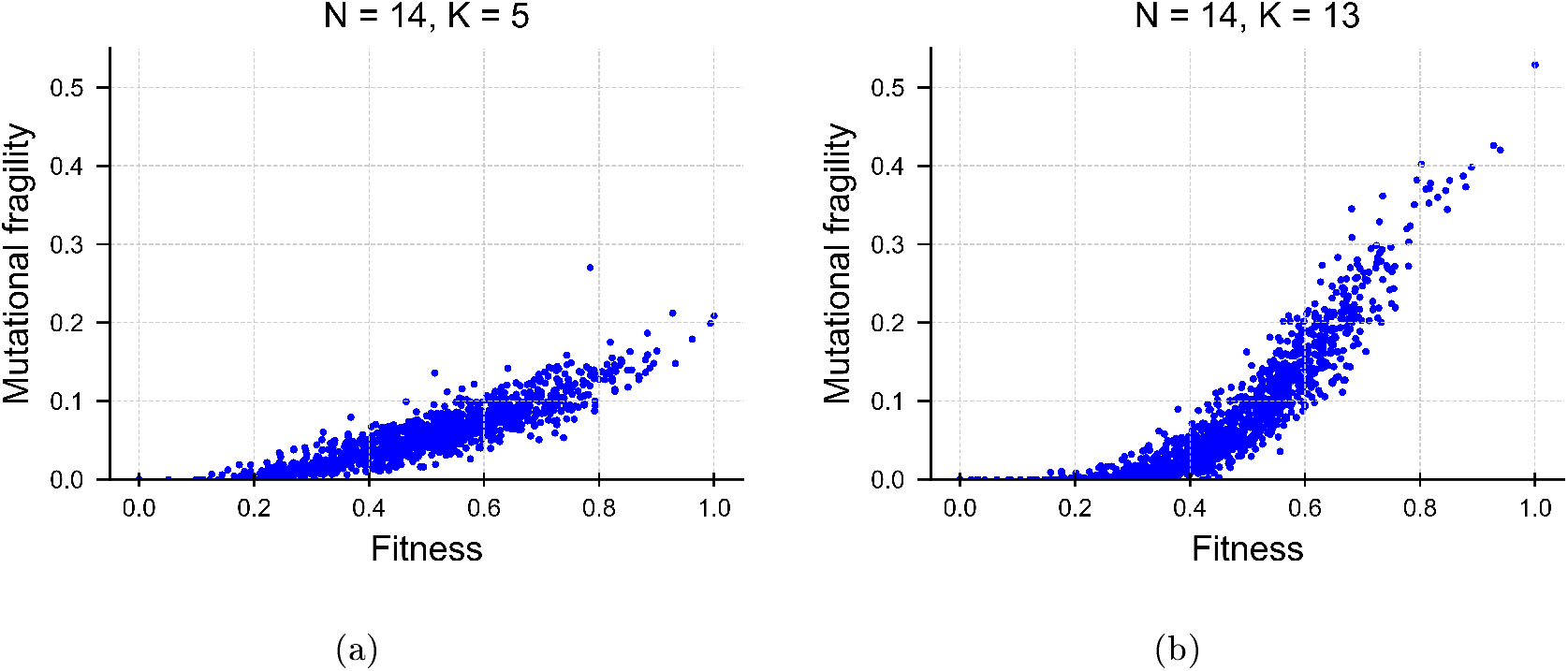
Mutational fragility in the NK model. Fragility vs. fitness scatter plots **(a-b)** in a simulated NK landscape with parameter values *N* = 14 *K* = 5 (a) and *K* = 13 (b). Low-fitness genotypes have mostly beneficial single-mutants and hence low mutational fragility (zero if all mutants are beneficial). The higher the fitness of a genotype, the more deleterious its single-mutants are, and hence it is more fragile. This effect is more pronounced in the uncorrelated landscape (*K* = *N* – 1 = 13, (b)) but is still observed in the partially correlated landscape with *K* = 5

## Discussion

Recent advances in high-throughput experimental methods have allowed for large-scale characterization of empirical fitness landscapes [4, 5, 6, 7], which can be applied to test hypotheses about the driving forces of evolutionary dynamics. Here we found a wild type which is sub-optimal and ruled out the possibilities that it is the fittest on average on multiple condition or located on a local fitness maximum. Instead, we found that the wild type is amongst the least mutationally fragile genotypes in the dataset, but the evolutionary mechanism at play here remains elusive. Robust (flat) genotypes were predicted theoretically [21, 22, 23], but were previously reported only in organisms having high mutation rates such as viruses [24, 25] or in digital organisms [26]. To the best of our knowledge, this is the first report of a flat gene in a low-mutation rate organism. It is increasingly appreciated that mutation rates are non-uniform across the genome and specifically, tRNA genes were shown to have 7-10 fold higher mutation rates compared to the background genome [27, 28]. Yet, it is unclear whether such mutation rate is sufficient to select for flatness. As the number of large-scale fitness measurements of particular landscapes is still limited, additional examples for wild type genes being sub-optimal are scarce, but see [29, 30].

Inherent to the astronomical dimensionality of fitness landscapes is our inability to fully measure them. Full landscape mappings are possible only for computationally fabricated landscapes [8, 31, 32]. Thus, usage of fragments of a fitness landscapes to draw general conclusions is a common practice in the field. It does raise the fundamental question whether indeed it represents the entirety of the landscape and hence, should be used with caution. Recent works handled the sparse sampling of fitness datasets by interpolating between the measured points to estimate fitness values of missing genotypes [33, 34] with some success. While these techniques are computationally very demanding, it would be interesting to test in the future whether they are applicable for computing evolutionary dynamics on incomplete landscapes.

## Acknowledgements

We thank Chuan Li and Jianzhi Zhang for sharing their experimental data and Ohad-Noy Feldheim for help with the fragility calculation. We thank Yoav Ram and Daniel Weissman for comments on an earlier version of the manuscript. T.F. acknowledges funding from the Hebrew University of Jerusalem. Y. P. acknowledges grant support from the Minerva foundation.

## Data availability

Upon submission we will deposit all the relevant resources in a public repository.

