## Supplementary methods, Supplementary text for "Fitness landscape analysis of a tRNA gene reveals that the wild type allele is sub-optimal, yet mutationally robust"

May 1, 2022

### 1 Methods

#### 1.1 Fitness data

We used the fitness measurements as published in [1]. For completeness, we briefly summarize how fitness was measured and defined there:

1. Cells were sampled and sequenced at time $T_0$, right before competition. The frequencies of the different genotypes $x_i(0)$ were then calculated $x_i(0) = \frac{R_i(0)}{\sum_j R_j(0)}$, where $R_j(t)$ is the number of reads of genotype $j$ at time $t$. These baseline frequencies were then used for all conditions.

2. The original cell pool was then split to the four different conditions, with at least 3 replicates for each (30°C and 23°C were replicated 6 times, the rest 3 times).

3. All genetic variants were grown together for 24 h, where after 12 h, the culture was diluted by a factor of 1/100.

4. After 24 h (T24) cells from each of the growth conditions were sampled and sequenced.

The number of wild type generations in 24 h under condition $m$ during competition was calculated as:

$$G^m_{\mathrm{WT}} = \log_2 \left( d \cdot \frac{g^m(24) x^m_{\mathrm{WT}}(24)}{g(0) x_{\mathrm{WT}}(0)} \right), \tag{S1}$$

where $g^m(t)$ is the total number of cells at time $t$ (calculated using the culture cell density (OD) measure) at condition $m$ and $d$ is the dilution factor. The measurements at time $t = 0$ were common to all conditions. The per-generation fitness of variant $i$ at condition $m$ was defined there as

$$f^m_i = \left( \frac{R^m_i(24)/R_i(0)}{R^m_{\mathrm{WT}}(24)/R_{\mathrm{WT}}(0)} \right)^{1/G^m_{\mathrm{WT}}(24)}. \tag{S2}$$

$G^m_{\mathrm{WT}}$ is the number of wild type generations reached after 24 h under condition $m$, $G^m_{\mathrm{WT}}(t) = r^m_{\mathrm{WT}} \cdot t/\log 2$, where $r^m_{\mathrm{WT}}$ is the wild type exponential growth rate under that condition. In the following we refer to $t = 24h$, and omit the time for brevity. Turning to continuous time and assuming that all variants grow exponentially during the entire experiment (neglecting lag and yield phases), Eq. (S2) can be written as

$$f^m_i = \left( \frac{\exp(r^m_i \cdot t)}{\exp(r^m_{\mathrm{WT}} \cdot t)} \right)^{1/G^m_{\mathrm{WT}}} = \exp\left[ (r^m_i - r^m_{\mathrm{WT}}) t \cdot \frac{\log 2}{r^m_{\mathrm{WT}} t} \right] = 2^{\left( \frac{r^m_i}{r^m_{\mathrm{WT}}} - 1 \right)} = 2^{\left( \frac{G^m_i}{G^m_{\mathrm{WT}}} - 1 \right)}, \tag{S3}$$

where $r_i^m$ is the growth rate of the $i-th$ genotype under condition $m$. Hence, the fitness of a genotype $i$ under condition $m$ is its exponentiated relative growth rate difference with respect to the wild type's under that condition. In the common notation of population genetics, a mutant has fitness advantage $s$ over the wild type, if it has on average $(1 + s)$-times more offspring per-generation. Thus,

$$f_i = 2^{\Delta r / r_{\mathrm{WT}}} = f_{\mathrm{WT}} \cdot (1 + s) = 1 + s, \tag{S4}$$

as the wild type's fitness equals 1 by definition, under each of the conditions. Hence,

$$s = 2^{\Delta r / r_{\mathrm{WT}}} - 1. \tag{S5}$$

#### 1.2 Fitness value re-scaling between conditions

The fitness values of genotypes were defined relative to the wild type's under each of the conditions and thus are incomparable between conditions. In order to calculate geometric mean fitness over all conditions, we first needed to define a common baseline to compare values referring to different conditions. Fitness of genotype $i$ under condition $m_2$ relative to the wild type's fitness at $m_2$ was defined as:

$$f_i^{m_2} := 2^{\left(\frac{r_i^{m_2}}{r_{\mathrm{WT}}^{m_2}} - 1\right)}. \tag{S6}$$

Now we would like to define it when the reference is the wild type's fitness at condition $m_1$, namely:

$$\tilde{f}_i^{m_2} := 2^{\left(\frac{r_i^{m_2}}{r_{\mathrm{WT}}^{m_1}} - 1\right)}. \tag{S7}$$

Substituting Eq. (S6) we obtain

$$r_i^{m_2} = \log_2(f_i^{m_2}) \cdot r_{\mathrm{WT}}^{m_2} + r_{\mathrm{WT}}^{m_2} = r_{\mathrm{WT}}^{m_2}(1 + \log_2(f_i^{m_2})). \tag{S8}$$

Substituting $r_i^{m_2}$ into Eq. (S7) we obtain:

$$\tilde{f}_i^{m_2} = 2^{(\log_2(f_i^{m_2}) \cdot r_{\mathrm{WT}}^{m_2} + r_{\mathrm{WT}}^{m_2} - r_{\mathrm{WT}}^{m_1}) / r_{\mathrm{WT}}^{m_1}}. \tag{S9}$$

If we return to the original reference growth rate $r_{\mathrm{WT}}^{m_2}$, the equation reduces to the original fitness definition, such that $\tilde{f}_i^{m_2} = f_i^{m_2}$.

We chose the measurements at 30°C to be our reference. For this calculation, we used the wild type growth rates under the four conditions as reported in [1]: $r_{\mathrm{WT}}^{23\mathrm{C}} = 0.25$, $r_{\mathrm{WT}}^{30\mathrm{C}} = 0.5$, $r_{\mathrm{WT}}^{\mathrm{DMSO}} = 0.45$, $r_{\mathrm{WT}}^{37\mathrm{C}} = 0.43$. The growth rate units are $\Delta$ OD/hour.

#### 1.3 Fragility calculation

Our basic definition of the mutational fragility of genotype $i$ is the average fitness difference between the genotype and its single mutants $N_1(i)$, where only deleterious mutants are accounted for while beneficial mutants are weighted as zero:

$$s_i = \frac{1}{|N_1(i)|} \sum_{j \in N_1(i)} |f_i - f_j|_+ \tag{S10}$$

$$|x|_+ = \begin{cases} x & \text{if } x \geqslant 0 \\ 0 & \text{if } x < 0 \end{cases}$$

where $f_j$ are the fitness values of these single mutants and $|N_1(i)|$ is the total number of single mutants (both deleterious and beneficial ones). This measure captures the sensitivity of genotypes to the harmful effect of mutations. However, our dataset contains fitness values of only a small subset of the tRNA gene fitness landscape with non-uniform sampling of the genotype space: dense close to the wild type and sparser further away. Only for the wild type, we have nearly full coverage of its single mutants, whereas for most other genotypes, only a few of their single mutants were measured. Moreover, for most genotypes under consideration the fitness values of the few single mutants that are closer or equally distant from the wild type are usually known. In contrast, for most genotypes, only few of the many single mutants that are further away from the wild type are known. This non-uniform sampling could potentially bias the fragility calculation.

To handle this, we weighted the fitness differences that are known such that they represent the proper size of their distance-from-wild type group. For a gene of length $L$, every genotype has $3L$ different single point mutations, as each of the $L$ positions can be mutated to 3 different bases. For a genotype at Hamming distance $d$ from the wild type, $d$ single mutants are closer to the wild type, $2d$ are equally distant from the wild type and the remaining $3L - 3d$ are further away. The weighted fragility measure for genotype $i$ of length $L$ at distance $d$ from the wild type then reads:

$$\begin{aligned}\hat{s}_{i,d(i)} = & \frac{d(i)}{3L}\frac{1}{|N_1(i)|}\sum_{j\in N_1(i),d(j)<d(i)}|f_i - f_j|_+ \\ & + \frac{2d(i)}{3L}\frac{1}{|N_1(i)|}\sum_{j\in N_1(i),d(j)=d(i)}|f_i - f_j|_+ \\ & + \frac{3L-3d(i)}{3L}\frac{1}{|N_1(i)|}\sum_{j\in N_1(i),d(j)>d(i)}|f_i - f_j|_+ .\end{aligned} \tag{S11}$$

The tRNA gene was mutated at 69 positions and hence each genotype can have up to 207 single mutants ($3 \times 69 = 207$). For example, consider a genotype that is a double mutant of the wild type. Out of the total 207 mutants, 2 single mutants are at Hamming distance 1 from the wild type (one of the two mutations is reverted) and 4 are at distance 2 (one of the already mutated positions is mutated to a different base). The remaining 201 mutants are at Hamming distance 3 from the wild type (additional mutation at a previously non-mutated position). In the fragility calculation we distinguish the 3 groups of mutants and weigh the fitness difference of the distance-1 mutants by 2/207, the distance-2 mutants by 4/207 and the distance-3 mutants by 201/207.

We included in the calculation only genotypes that had at least 5 single mutants that were measured and for which at least 3 of them are further from the wild type than the genotype under consideration.

#### 1.4 Simulated fitness landscape using the NK model

We simulated a correlated fitness landscape using the NK model [2]. A genotype in this model is represented by a binary string of length $N$, such that the fitness landscape consists of $2^N$ genotypes. The parameter $K$ is used to tune the ruggedness of the fitness landscape, such that for $K = 0$ it is fully additive and smooth and for $K = N - 1$ it is the most rugged. The fitness of each genotype is defined as the average of $N$ contributions of its $N$ positions. Each of these contributions is determined by the sequence content at $K+1$ positions: the one under consideration and $K$ other positions, with which it supposedly interacts. We drew $2^{K+1}$ random numbers from a uniform distribution and assigned these values to be the fitness contributions of all the possible binary strings of length $K+1$, $f(s_0, \ldots, s_K)$, $s_i \in [0, 1]$. The fitness of each genotype encoded by a binary string of length $N$ is defined as the average of the $N$ fitness contributions of the length-$K+1$ strings it contains (cyclically): $F(s_0, s_1, \ldots, s_{N-1}) = \frac{1}{N} \sum_j f(s_j, s_{j+1}, \ldots, s_{(j+K) \mathrm{mod} N})$.

### 2 Assessment of fitness measurement errors

The fitness values were calculated in [1] by sequencing culture samples at two time points, $T = 0$, $T = 24$ h and estimating the relative frequencies of the different genotypes in the culture. Fitness calculation assumed that the cells grew exponentially between these two time points (See above). Our focus in this paper was on the fraction of genotypes with fitness higher than the wild type's. Below, we scrutinize potential sources of error in the fitness value estimate, in order to rule out the possibility that these high-fitness values are due to an experimental artifact.

#### 2.1 Exponential growth assumption

The underlying assumption in the fitness calculation is that all genotypes grew exponentially during the entire experiment (24 h.). In practice, before entering exponential growth, cells spend time in a preparatory phase called a 'lag phase' (in which growth rate is relatively slow). The length of the lag phase could vary between strains, hence sampling all genotypes at the same time could potentially catch some of them still in the lag phase. In a batch culture, once the cells use up the available nutrients, their numbers saturate and growth slows down again (also called 'yield' phase). Although the culture was diluted after 12 h [1] some strains could have reached their yield phase earlier than others. As we do not have detailed growth curves, it is hard to assess how common this is and to what extent it could have affected the calculated fitness values. However, this error source, if it exists, is expected to only *underestimate* fitness values. That is because, the final genotype frequency could have been attained in a shorter time period than assumed in the calculation, and thus the growth rate must have been larger than estimated.

#### 2.2 Read-count noise and variation between biological repeats

The frequencies of the different genotypes in the culture were estimated by Li *et al.* using samples of the batch culture at two time points where only genotypes with at least 100 reads at $T = 0$ were considered [3]. Sampling errors could potentially lead to error in fitness estimates. In contrast to the previous error source, which could only cause under-estimation of fitness values, read-count noise could cause both over and under-estimation of fitness. For example, a combination of under-sampling at $T = 0$ and over-sampling at $T = 24$ h of a particular genotype could cause an *over-estimate* of its fitness. In contrast, over-sampling at $T = 0$ and under-sampling at $T = 24$ h should lead to the opposite effect of fitness value *under-*

*estimation.* Sampling errors in $T = 0$ and $T = 24$ h could also (at least partially) cancel each other, if they are both in the same direction.

In the fitness assessment protocol by Li and co-workers, a common sampling of all genotypes at $T = 0$ was applied [3]. This common culture was then used to inoculate the multiple growth experiments under the four different growth conditions, with 5 (23°C, 30°C) or 3 (37°C, oxidative stress) independent replicates under each condition. Thus, at $T = 24$ h, there were multiple samples for each genotype under each condition (one for each replicate).

To estimate the effect of sampling errors on fitness, we used the raw data of the number of reads obtained in the experiments for each genotype, as provided by Li and co-workers. To estimate the distribution of fitness values, we assumed that the number of reads of genotype $i$ both at $T = 0$ and $T = 24$ are drawn from Binomial distributions $\text{Binom}(N, p_i)$, where $N$ is the total number of reads of all genotypes at the relevant time. We estimate $p_i$ by $\hat{p_i} = \frac{\text{Number of reads of genotype i}}{N}$ [4]. We then randomly drew 20 realizations of these distributions at these two time points. We calculated the different fitness values obtained by all possible combinations (total of 400) of the fabricated numbers of reads at $T = 0$ h and $T = 24$ h using Eq. (S2). We assumed that the number of reads of the wild type is fixed.

In Fig. S1a-b, we show these distributions of fitness at 30°C for two different genotypes: one with fitness close to the wild type's and one with very high fitness. We repeated this procedure for the five biological repeats, shown there using different colors. We find, that the variation incurred by read count in a single repeat is typically smaller than the variation between repeats. Thus, in the following we focus on the between-repeat variation as the dominant error source.

The fitness values reported by Li *et al.* which we used in our calculations were averages of the fitness values obtained in the different biological repeats. Hence, the relevant error measure is the error of these average values. In Fig. S2 we show scatter plots of the relative standard error of the mean ('sem', standard deviation between repeats divided by square root of number of repeats) against the mean. One can appreciate that the relative sem is smaller than 0.1 in all cases and for most genotypes smaller than 0.05. This rules out the possibility that the $\sim$2000 fitter than wild type genotypes are considered as such only due to measurement inaccuracy.

#### 2.3 Background mutations

Mutations could have also occurred in other genome locations during the course of the competition experiment. While not a measurement error, had such a mutation occurred, we could have mistakenly attributed the fitness effect to the tRNA variant. The cells barcode identifies them by their tRNA variant. If such a background mutation occurred during the course of the experiments, after cells had already started reproducing, it would result in cells carrying a common barcode that are a mixture of multiple sub-populations: the original genome and the one carrying the background mutations. The later such a mutation emerges, the smaller its population fraction. Since sequencing cannot distinguish between the two types, the estimated fitness would average over the two sub-populations. Thus, the effect of the background mutation would be largest, if it occurred at the very beginning when only a single cell of each tRNA variant existed. We are particularly interested in determining whether high-fitness genetic variants can be explained by such background mutations; hence below we will focus

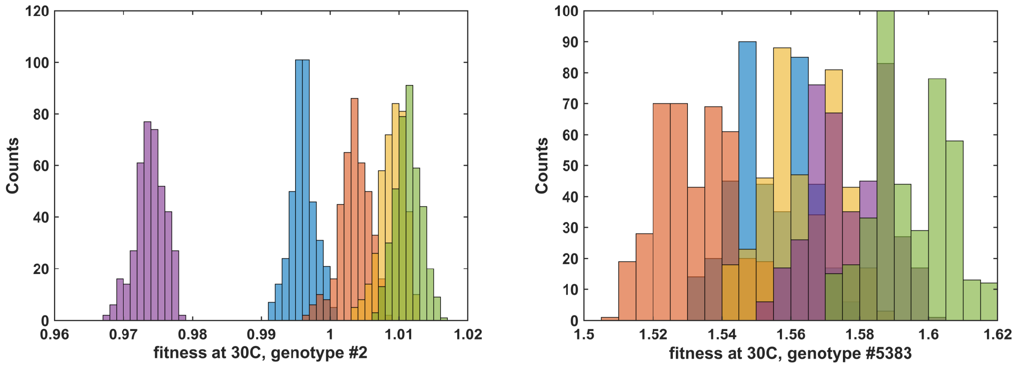


(a) (b)

Figure S1: **Estimation of potential errors in fitness values due to read-count inaccuracy.** We show here the fitness values distributions for the different genotypes, one with fitness close to the wild type's **(a)** and one with highest fitness **(b)**. These fitness distributions were calculated by randomly drawing realizations from the read count distributions at both $T = 0$ h and $T = 24$ h, assuming they are Binomial. For each genotype we independently assessed the fitness distributions obtained at each biological repeat. They are shown here using a different color for each repeat. We observe that the variation between biological repeats is at least comparable to the read count noise **(b)** if not larger **(a)**.

on beneficial mutations alone. Both neutral and deleterious mutations are expected to have only a minor effect on fitness estimates. Neutral mutations have no effect on fitness to begin with and deleterious mutations are naturally selected against and hence will be present in low numbers.

In the experimental procedure used by Li and co-workers, the different variants were synthesized using error-prone PCR initiated with the wild type strain. The products were then transformed into yeast cells (See [1, 3] SI for more details). We estimate that only a small fraction ($\approx 0.001\%$) of the yeast cells was transformed with a variant, and hence each variant exists in very few cells. For simplicity, we assume here that each variant exists as a single copy only, which is the worst-case scenario for our purpose.

To estimate the likelihood of naturally occurring beneficial mutations, we use results from Levy *et al.* [4], where the evolutionary dynamics of 500,000 different yeast lineages, each tagged by a unique barcode, was tracked. They reported the rate of beneficial mutations as a function of their fitness effect. For example, the rate of beneficial mutations that led to fitness increment $s > 5\%$ was approximately $10^{-6}$ per cell per generation. This is equivalent, in our formalism, to fitness $f_i = f_{\mathrm{WT}} \cdot (1 + s) = 1 + s > 1.05$. The expected number of such beneficial mutations that occurred in the first generation in any of the 23,284 different genotypes is only 0.02. If we account for mutations occurring during the entire transformation phase (which lasted 50 generations), the expected number of such beneficial mutations per generation increases, because the total number of cells increases. However, the cells having beneficial mutations are mixed with non-mutated cells carrying the same barcode. Hence, the fitness effect of the mutations needs to be much larger for it to have the same measured fitness effect when averaged with non-mutated cells. Since the spectrum of beneficial mutation rates reported by

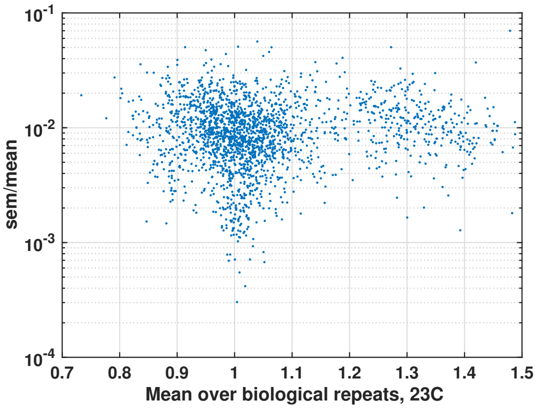


(a)

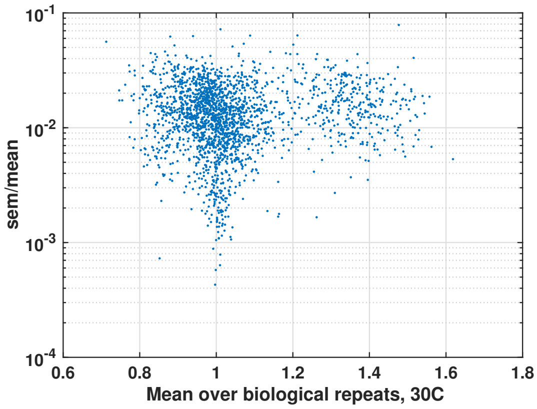


(b)

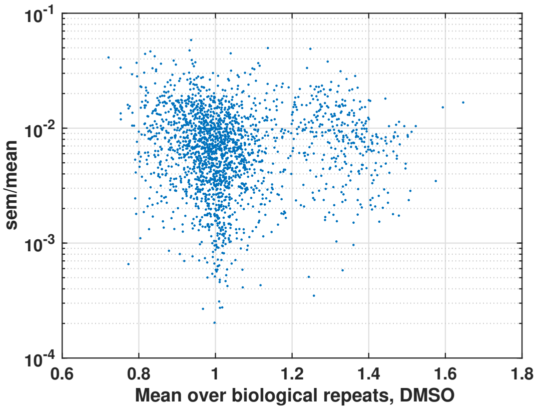


(c)

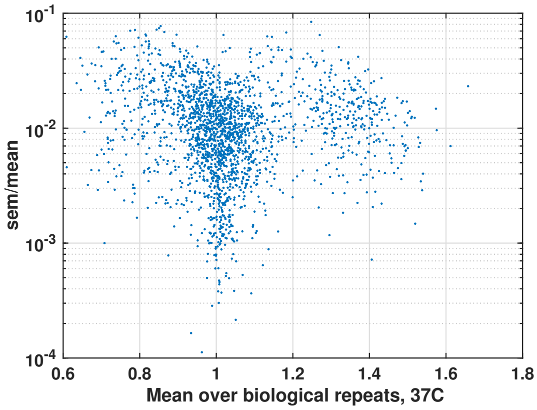


(d)

Figure S2: **Fitness variation between biological repeats under each condition.** We show here a scatter plot of the standard error of the mean calculated over 5 (23°C, 30°C) or 3 (37°C, oxidative stress) independent repeats divided by the mean and plotted against the reported fitness. We calculated this for all genotypes that had at least 50 reads in all experiments both at $T = 0$ and at $T = 24$. Every point represents a particular variant. Each panel shows one of the four conditions.

Levy *et al.* drops sharply near $s = 12\%$ fitness benefit, we estimate such a beneficial mutation to be far less likely. This leaves the expected number of beneficial background mutations during transformation to be dominated by mutations at the first generation, namely 0.02, far less from the reported ~2000 fitter than wild type variants found by Li *et al.* . Hence, this error source is obviously negligible.

#### 3 Mutational fragility in the NK model

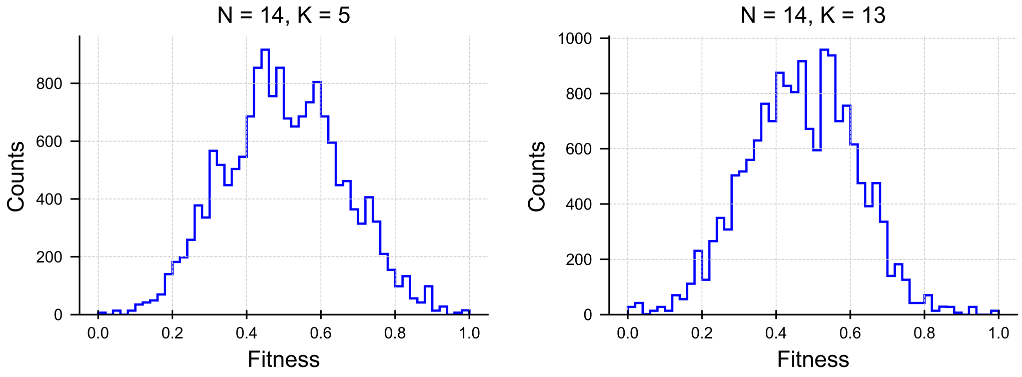


(a) (b)

Figure S3: **Fitness value distributions in NK model.** Fitness distributions in the simulated NK landscape with parameter values $N = 14$, $K = 5$ (a) and $K = 13$ (b). These simulated landscapes were used to produce the fragility-fitness scatter plots of Fig. 3 in the main text.
